# Evaluation of therapeutic effect of baloxavir marboxil against high pathogenicity avian influenza virus infection in duck model

**DOI:** 10.1101/2025.10.24.684283

**Authors:** Yo Shimazu, Norikazu Isoda, Takahiro Hiono, Hew Yik Lim, Daiki Kobayashi, Misato Shibazaki, Nguyen Bao Linh, Tatsuru Morita, Ryo Daniel Obara, Mariko Miki, Hiromi Osaki, Takao Shishido, Yoshinori Ikenaka, Yoshihiro Sakoda

## Abstract

Since 2020, high pathogenicity avian influenza virus (HPAIV) infections in wild birds have been frequently reported. Because HPAIV infection has occasionally caused outbreaks in captive rare birds, application of antiviral drugs for treatment purposes against them has been considered from the perspective of conservation medicine. In this study, the therapeutic efficacy of baloxavir marboxil (BXM) was evaluated using a duck model to help establish the post-infection treatment for rare birds. Sixteen four-week-old ducks were divided into four groups and intranasally inoculated with the HPAIV strain A/crow/Hokkaido/0103B065/2022 (H5N1). BXM was orally administered once daily at doses of 12.5, 2.5, 0.5, and 0 mg/kg to each of the four groups from 2 to 6 days post-infection. Blood samples were collected at 2, 8, and 24 hours after the initial BXM administration to measure the plasma concentrations of its active form, baloxavir acid (BXA). All ducks were monitored until 14 days post-infection, and their oral and cloacal swabs were collected for virus recovery. All eight ducks administered with 12.5 or 2.5 mg/kg of BXM survived, demonstrating a significant reduction in virus recovery compared to the 0 mg/kg group. Pharmacokinetic/pharmacodynamic (PK/PD) analysis of BXA suggested that parameters such as C_max_ and AUC_0–24hr_ were correlated with the suppression of virus shedding. These findings demonstrated that BXM administration within 48 hours post-HPAIV infection in ducks effectively reduced mortality and virus shedding. The comparison of PK parameters may help estimate efficient BXM dosing strategies in rare birds.

**Highlight:** BXM prevented death of HPAIV-infected ducks when given within 48 hr of inoculation.

Doses of 2.5 mg/kg rapidly suppressed and eliminated infectious virus shedding.

AUC_0–24hr_ and C_max_ correlated with suppression of virus shedding.

## 1. Introduction

High pathogenicity avian influenza viruses (HPAIVs), classified as members of influenza A viruses (*Alphainfluenzavirus influenzae*), cause fatal diseases in various bird species (Alexander and Brown, 2009). In recent years, these viruses have spread widely among wild bird populations, and their impact is now recognized on a global scale (Xie et al., 2025). Since 2020, HPAIV infections have been continuously reported in various regions, including Europe, Asia, Africa, America, and Antarctica (Banyard et al., 2024; Sacristán et al., 2024). These epidemics have been shifting from a seasonal to a year-round endemic situation in Europe. Under these circumstances, HPAIV outbreaks have also occurred among captive birds in zoos, resulting in the death of rare bird species (Schlachter et al., 2025; Usui et al., 2020). While vaccination has been implemented in some zoos during domestic outbreaks of HPAIVs in several countries, such as Singapore, the Netherlands, Spain, Denmark, and the United States (Bertelsen et al., 2007; Katzner et al., 2025; Oh et al., 2005; Philippa et al., 2005; Vergara-Alert et al., 2011), the use of vaccines in captive birds remains unavailable in several countries, including Japan. In these countries, infection control relies mainly on enhanced biosecurity measures, and once HPAIV infection is detected in captive birds, infected and contact birds are recommended to be culled to prevent further spreading of infection and eliminate the virus (Liu et al., 2020). Nevertheless, from the perspective of conservation medicine, therapeutic options for rare birds should be explored. In zoos, birds are often housed in open cages from the perspective of animal welfare, leading to limitations in biosecurity. Under these circumstances, not only prophylactic but also therapeutic countermeasures are important. Currently, there is no established treatment protocol for HPAIV infection in captive birds, and the development of effective antiviral treatment is urgently needed.

For human influenza virus infection, several antiviral drugs targeting the viral neuraminidase (NA) and the polymerase acidic protein (PA) are widely used. The NA, a surface glycoprotein of the influenza virus, exhibits sialidase activity and facilitates virus release from infected cells (Itzstein et al., 1993). The NA inhibitors oseltamivir, zanamivir, and peramivir have been evaluated for their antiviral effects against HPAIVs in a chicken model (Meijer et al., 2004; Twabela et al., 2020). However, none of these drugs completely prevented the onset of disease and death. Even when administered simultaneously with HPAIV challenge, they reduced mortality by only about 80% (oseltamivir), 0% (zanamivir), and 75% (peramivir), respectively. NA inhibitors suppress viral release in the late stages of infection, but since they do not inhibit viral replication, their effect is thought to be limited. The PA, a subunit of the viral RNA-dependent RNA polymerase complex of the influenza virus, is involved in cap-snatching, acquiring capped RNA fragments from host mRNAs via its endonuclease activity (Dias et al., 2009). PA inhibitors show higher antiviral efficacy than NA inhibitors by suppressing viral replication in the early stages of infection (Checkmahomed et al., 2020). Baloxavir marboxil (BXM) is a PA inhibitor that has been approved for clinical use in humans, but not in animals. Simultaneous administration of BXM at 2.5 mg/kg or higher to chickens inoculated with HPAIV indicated its prophylactic effect of disease onset suppression (Twabela et al., 2020). Assuming an initial infection case in zoos, treatment would be initiated after captive birds had shown clinical symptoms and been subjected to diagnosis. The previous study also evaluated the efficacy of BXM following post-infection administration in a chicken model; however, even administering 20 mg/kg of BXM with 12-hour intervals from 24 hours post-infection, all individuals died, and the efficacy of post-infection administration was not observed (Twabela et al., 2020). This result is likely due to the extremely rapid disease progression of HPAIVs in chickens, which typically results in death within 2–3 days post-infection (Spickler et al., 2008), thus heavily limiting the time window for therapy in the chicken model. However, infection with HPAIVs in some non-Galliform species, such as sparrows, crows, herons, white-tailed eagles, and wild waterfowls, often resulted in slower progression (4–7 days from infection to death) or was even sublethal (Bordes et al., 2024; Fujimoto et al., 2022; Hiono et al., 2016; Soda et al., 2022). Therefore, an avian model with slower disease progression is needed for evaluating the post-infection efficacy of BXM administration to develop treatment strategies applicable to rare bird species.

In this study, the duck was utilized as a model animal, because it is considered more suitable for extrapolating the outcome to rare bird species based on its slower disease symptom progression compared to the chicken model (Bordes et al., 2024). BXM was administered to infected ducks, and their survival rates, clinical symptoms, and virus shedding were monitored over time. Based on the results, the effective dosage and optimal schedule of BXM administration were suggested. Furthermore, pharmacokinetic parameters in plasma samples collected after BXM administration were analyzed to investigate their association with antiviral effects. The outcome of this study will provide a fundamental reference for establishing practical treatment strategies against HPAIV infection in rare bird species.

## 2. Materials and methods

### 2.1. Viruses

Three HPAIVs, A/crow/Hokkaido/0103B065/2022 (H5N1) (Cr/Hok/22) (Isoda et al., 2022), A/black swan/Akita/1/2016 (H5N6) (Bs/Aki/16) (Okamatsu et al., 2017), and A/whooper swan/Hokkaido/4/2011 (H5N1) (Ws/Hok/11) (Sakoda et al., 2012), were isolated from a dead large-billed crow (*Corvus macrorhynchos*) found in an urban garden, a black swan (*Cygnus atratus*) bred at a zoo, and a wild whooper swan (*Cygnus cygnus*) found at the waterside of their resting areas, respectively. They were propagated in 10-day-old embryonated chicken eggs for 44 hours at 35°C. The collected allantoic fluids were stored as the virus stock for experimental infection.

### 2.2. Animals

Twenty-five four-week-old Cherry Valley ducks (*Anas platyrhynchos* var. *domesticus*, 1.4–2.1 kg body weight, Anshin-Seisan Farm, Horonobe, Hokkaido, Japan) were used in this study. Before starting the experiment, it was confirmed that ducks did not possess antibodies against the influenza A virus using the IDEXX Influenza A Ab Test (IDEXX Laboratories, Inc., Westbrook, ME, U.S.), an enzyme-linked immunosorbent assay (ELISA) kit.

### 2.3. Cells

Madin-Darby canine kidney (MDCK) cells were maintained in minimum essential medium (MEM) (Shimadzu Diagnostics Corp., Tokyo, Japan) supplemented with 0.3 mg/mL L-glutamine (FUJIFILM Wako Pure Chemical, Osaka, Japan), 5% fetal bovine serum (Thermo Fisher Scientific, Waltham, MA, USA), 100 U/mL penicillin G (Meiji Seika Pharma, Tokyo, Japan), 0.1 g/mL streptomycin (Meiji Seika Pharma), and 8.0 µg/mL gentamicin (MSD, Rahway, NJ, USA).

### 2.4. Experimental infection of ducks

As a preliminary test to select the virus strain for the next experiment, nine ducks were divided into three groups and challenged with 10^6.0^ times of 50% egg infectious dose (EID_50_) of Cr/Hok/22, Bs/Aki/16, and Ws/Hok/11, respectively. Clinical scores were determined by applying the criteria used for the calculation of the intravenous pathogenicity index (IVPI) according to the World Organisation for Animal Health (WOAH) manual (WOAH, 2024). Briefly, birds were scored 0 if healthy, 1 if sick, 2 if severely sick, and 3 if dead. Birds were considered sick if one of the following signs was observed, and severely sick if more than one of the following signs was observed: respiratory involvement, depression, diarrhea, cyanosis of the feet or mucosa, edema of the face or head, and nervous signs. When the ducks were unable to drink or eat, they were deemed to have reached the humane endpoint and euthanized. From the following day, they were scored as 3.

Sixteen ducks were divided into four groups, and all ducks were intranasally challenged with 10^6.0^ EID_50_ of Cr/Hok/22. BXM (20 mg tablets, Shionogi, Osaka, Japan) was ground and suspended in saline (Otsuka Pharmaceutical Factory, Inc., Tokushima, Japan). BXM suspension was administered orally to three groups of ducks once daily at the doses of 12.5, 2.5, or 0.5 mg/kg, respectively, from 2 to 6 days after the HPAIV challenge. Saline was administered to the 0 mg/kg group during the study period. The five-day administration regimen was designed considering the shorter elimination half-life of BXA in ducks compared with humans, to maintain adequate plasma exposure during treatment (Twabela et al., 2020). Clinical manifestations were observed daily. Oral and cloacal swabs of the challenged birds were collected at 0–7, 9, 11, 14 days post-infection (dpi) and suspended in 2 mL of viral transport medium consisting of MEM (Shimadzu Diagnostics) containing 10,000 U/mL of penicillin G (Meiji Seika Pharma), 10 mg/mL streptomycin (Meiji Seika Pharma), 0.3 mg/mL gentamicin (MSD), 0.05 mg/mL nystatin (Meiji Seika Pharma), and 0.5% of bovine serum albumin fraction V (Roche Diagnostics, Mannheim, Germany). Viral titers were calculated by the method of Reed and Muench (Reed and Muench, 1938) and expressed as 50% tissue culture infectious dose (TCID_50_) per milliliter of swab suspension.

In order to measure the plasma concentration of baloxavir acid (BXA), the active form of BXM, 230 μL of blood samples were collected from ducks administered BXM of 12.5 and 2.5 mg/kg at 2, 8, and 24 hours post-initial dose (at 2, 3 dpi), before the fifth dose, and 2, 8, and 24 hours post-fifth dose (at 6, 7 dpi). The collected blood samples were mixed thoroughly with 13.8 μL of an enzyme inhibitor/anticoagulant mixture (60 mmol/L dichlorvos (FUJIFILM Wako Pure Chemical)/sodium heparin (Nipro Corp., Osaka, Japan)), and centrifuged at 1900 g for 15 minutes to obtain plasma.

All animal experiments were carried out at the animal BSL-3 facility at the Faculty of Veterinary Medicine, Hokkaido University, which has been accredited by the Association for Assessment and Accreditation of Laboratory Animal Care International since 2007, with approval from the Institutional Animal Care and Use Committee of the Faculty of Veterinary Medicine, Hokkaido University (approval number: 23-0052).

### 2.5. Quantitative RT-PCR

Viral RNAs were extracted from oral and cloacal swab samples using the QIAamp viral RNA mini Kit (Qiagen, Venlo, Netherlands) according to the manufacturer’s instructions and were stored at –80℃ until further processing. The presence of the matrix gene was investigated by quantitative reverse transcription-polymerase chain reaction (qRT-PCR) using by THUNDERBIRD Probe One-step qRT-PCR Kit (Toyobo, Osaka, Japan) on a LightCycler 480 System (Roche Diagnostics K.K., Tokyo, Japan) with the primer sets as described by Heine et al. 2015 (Heine et al., 2015).

### 2.6. Pharmacokinetic analysis

The plasma concentration of BXA was determined using a liquid chromatograph-tandem mass spectrometer (Agilent 6495B, Agilent Technologies, Inc., Santa Clara, CA, USA). The limit of detection of BXA was 0.004 µg/mL. The pharmacokinetic (PK) parameters of maximum plasma concentration (C_max_), area under the plasma concentration-time curve from 0 to 24 hours (AUC_0–24hr_), and elimination rate constant (k_el_) were determined using the linear trapezoidal method, and half-life in plasma (T_half_) was calculated from k_el_.

### 2.7. Pharmacokinetic (PK) / Pharmacodynamic (PD) analysis

To assess the relationship between PK parameters and antiviral efficacy, the independent variables (x-axis) were set as the PK parameters AUC_0–24hr_, C_max_, and plasma concentration of BXA at 24 hours post administration (C_24hr_), respectively. The dependent variable (y-axis) was defined as the difference in viral titers between 2- and 3-days post-infection. Logarithmic trend lines were fitted using Excel (Microsoft, Redmond, WA, U.S.), and the corresponding correlation coefficients (R² values) were calculated.

### 2.8. Statistical analysis

The mean and standard deviation (SD) of values for clinical score, virus titer, and blood plasma concentration of BXA were calculated using Excel (Microsoft). A one-sided Student’s t-test was performed to compare those values between the groups of each dose and the untreated group. For virus titers below the detection limit, a value of 10^0.8^ TCID_50_ was assigned for statistical calculation. The survival rate of birds between the groups was compared using a log-rank test using OriginPro 2025 (OriginLab Corp., Northampton, MA, U.S.). When the P-value was less than 0.05, the difference relative to the control group was regarded as statistically significant.

## 3. Results

### 3.1. Antiviral effects of BXM against high pathogenicity avian influenza virus infection in ducks

All birds inoculated with Cr/Hok/22 died within 11 dpi (Figure S1A). One bird died at 9 dpi and two birds survived among the ducks inoculated with Bs/Aki/16. All three birds inoculated with Ws/Hok/11 survived. Mean clinical scores were also relatively high in birds inoculated with Cr/Hok/22 (Figure S1B). In order to evaluate the efficacy of BXM with precision, Cr/Hok/22 was utilized in the following experiments due to its high pathogenicity to ducks.

Four ducks per group were intranasally inoculated with 10^6.0^ EID_50_ of Cr/Hok/22. In the 0 mg/kg group, all ducks died within 7 dpi (Figure 1A). In the 0.5 mg/kg treatment group, one out of four treated birds died on 5 dpi, and the other three birds survived for the experimental period, indicating an apparent improvement in survival rate (*p*=0.056). None of the birds administered at 12.5 or 2.5 mg/kg died after virus challenge, demonstrating complete suppression of mortality in these groups compared to the 0 mg/kg group (*p*=0.007). No clinical symptoms—or only transient symptoms followed by recovery—were observed in any of the BXM-treated ducks, except for one duck in the 0.5 mg/kg group that died on 5 dpi and one duck in the 12.5 mg/kg group that had leg paralysis until the end of the study period despite recovering from acute symptoms (Figure 1B, Table S1).

**Figure 1.**
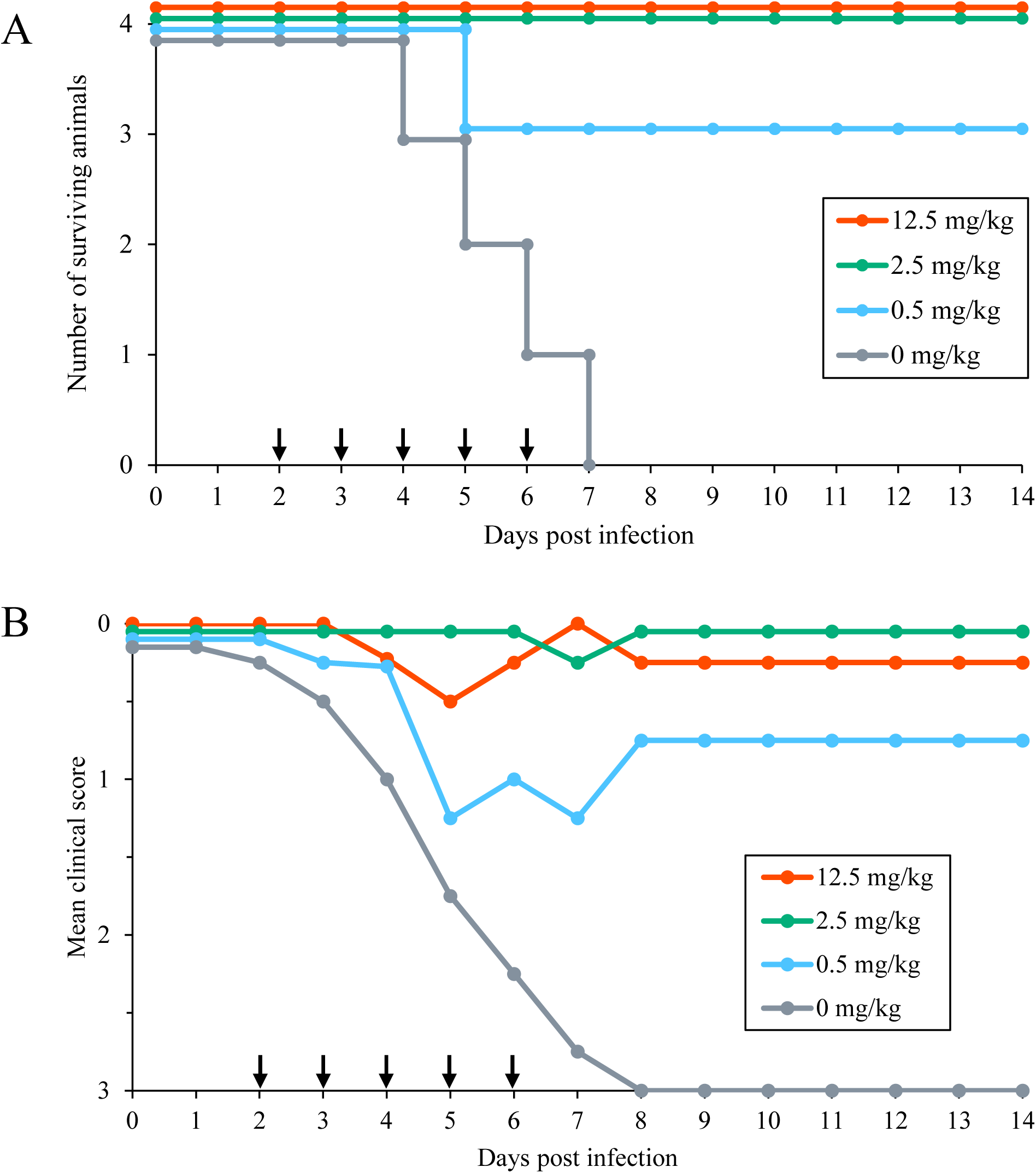
The efficacy of post-infection administration of baloxavir marboxil (BXM) in ducks infected with Cr/Hok/22. (A) Survival curves. (B) Mean clinical scores. BXM was administered orally at 12.5, 2.5, 0.5, or 0 mg/kg once daily on days 2–6 post-infection (arrows).

The virus titers recovered from oral swabs of the 0.5 mg/kg treated duck on 2 days post-initial dose (dpid) (equal to 4 dpi) were significantly lower than those of the 0 mg/kg group on that day (*p*=0.041) (Figure 2A). In the 12.5 and 2.5 mg/kg groups, the titers of virus recovered from oral swabs significantly decreased from 1 dpid (3 dpi) compared with the 0 mg/kg group (*p*<0.001 [12.5 mg/kg], p=0.0010 [2.5 mg/kg]), and were below the detection limit since 3 dpid (5 dpi). In cloacal swabs, the virus titers of the ducks treated with all three doses were significantly lower than those of the 0 mg/kg group at 1dpid (3 dpi) (p<0.001 [0.5 mg/kg], p=0.021 [2.5 mg/kg], p=0.001 [12.5 mg/kg]). Furthermore, since 2 dpid (4 dpi), all virus titers of the ducks administered with 12.5 and 2.5 mg/kg were below the detection limit of 10^0.8^ TCID_50_.

**Figure 2.**
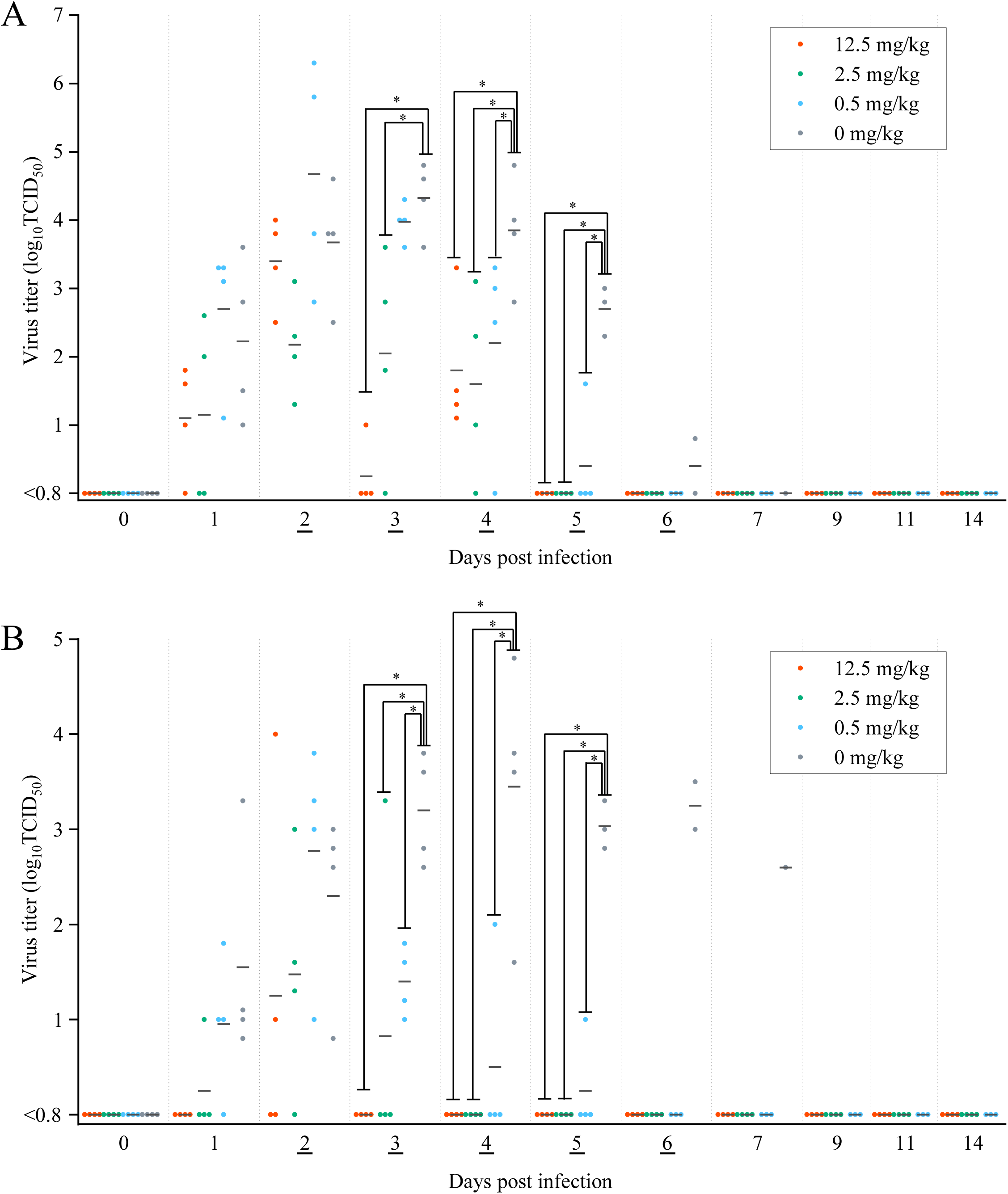
Virus titer of (A) oral and (B) cloacal samples in the ducks infected with Cr/Hok/22. BXM was administered orally at 12.5, 2.5, 0.5, or 0 mg/kg once daily on days 2–6 post-infection (underline). Bars represent the mean values. \**p*<0.05 versus 0 mg/kg administration group.

Although the infectious virus recovery from swab samples was limited to 2 or 3 dpid, viral genomes were continuously detected until 4, 5, and 5 dpid in oral swabs of the ducks administered with 12.5, 2.5, and 0.5 mg/kg of BXM, respectively (Table S2).

### 3.2. Pharmacokinetic analysis of baloxavir acid in ducks infected with high pathogenicity avian influenza virus

To evaluate the pharmacokinetics of BXA, the active form of BXM, plasma samples were collected at 2, 8, and 24 hours post-initial dose of BXM (2, 3 dpi) and at 0, 2, 8, and 24 hours post-fifth administration of BXM (6, 7 dpi), and plasma concentration of BXA in each sample was analyzed (Figure 3). After the initial administration of BXM at 12.5 or 2.5 mg/kg, the plasma concentration of BXA reached 1784 or 285 ng/mL at 2 hours post-administration, and decreased to 130 or 26 ng/mL at 24 hours post-administration, respectively. After the fifth administration of BXM, the plasma concentration of BXA increased to 876 or 145 ng/mL at 2 hours post-administration and decreased to 18 or 6 ng/mL at 24 hours post-administration, respectively. Based on these results, PK parameters were calculated. All parameters investigated for BXA plasma exposure, including AUC_0–24hr_, C_max_, and C_24hr_ increased dose-dependently. Additionally, AUC_0–24hr_, C_max_, and C_24hr_ of BXA after the fifth administration were lower compared to those observed after the initial administration during the 5-day repeated administrations.

**Figure 3.**
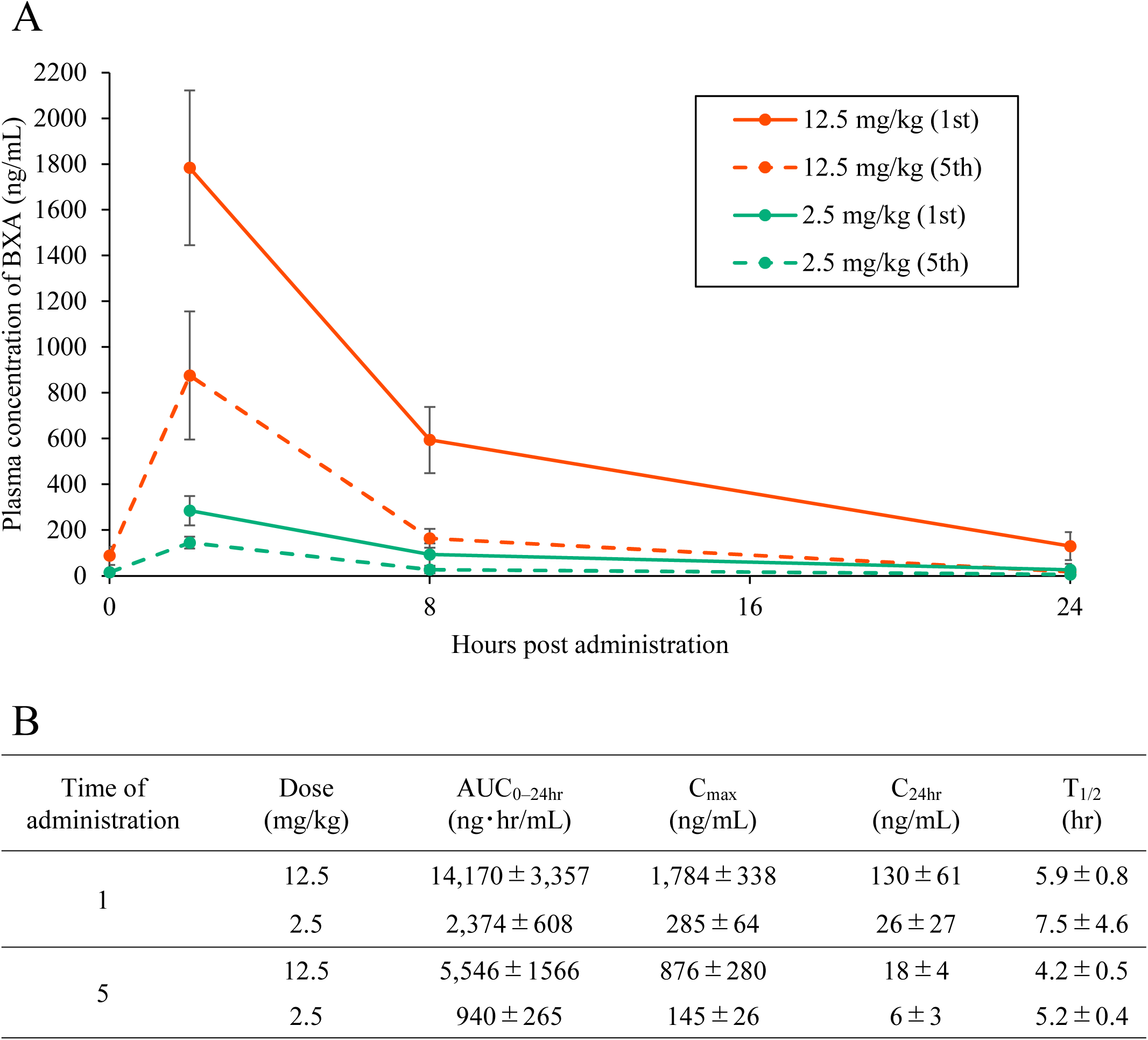
(A) Mean plasma concentration of BXA after the administration of BXM at 12.5 and 2.5 mg/kg. Data represent the mean±SD of four ducks. (B) Mean±SD pharmacokinetic parameters of BXA in the plasma in the ducks administered BXM at 12.5 or 2.5 mg/kg.

### 3.3. PK/PD analysis of baloxavir acid in ducks infected with high pathogenicity avian influenza virus

Correlations between PK parameters from the 12.5 and 2.5 mg/kg groups and changes in virus recovery before and after treatment were analyzed. In each type of swab, the difference in virus titers between the day of initial BXM administration (2 dpi) and the 1 dpid (3 dpi) was calculated. In oral swabs, reductions in virus titers correlated with both AUC_0–24hr_ and C_max_, with R^2^ values of 0.659 and 0.678, respectively. This suggested an association of these parameters with the inhibitory effect on virus shedding (Figure 4). In contrast, in cloacal swabs, C_24hr_ showed only a weak positive correlation with the reduction of virus shedding, with an R^2^ value of 0.333.

**Figure 4.**
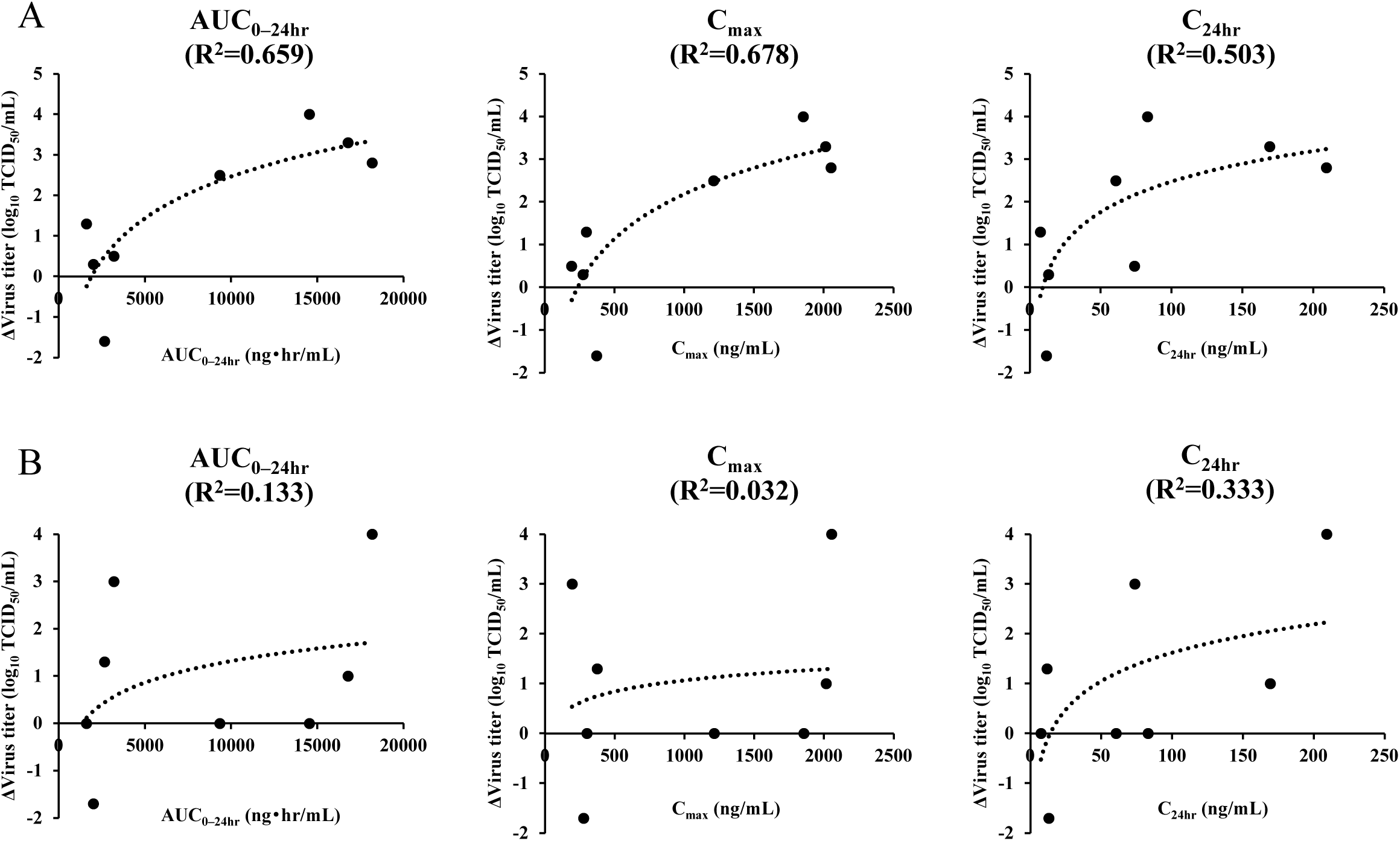
Estimated linear curves between each PK parameter and the reduction of virus titers in oral (A) and cloacal (B) swabs from 2 to 3 days post-infection.

## 4. Discussion

The application of human anti-influenza drugs against HPAIV infection in birds has been considered (Meijer et al., 2004; Twabela et al., 2020). While prophylactic administration of oseltamivir, zanamivir, and peramivir only partially reduced mortality in chickens infected with HPAIV, BXM suppressed the onset of HPAIV infection at a dose of 2.5 mg/kg or higher of simultaneous administration (Twabela et al., 2020). In the present study, ducks were utilized as they show typical disease manifestations for 4–7 days after HPAIV inoculation (Bordes et al., 2024), which is similar to that of HPAIV infection in wild birds. The results of the present study demonstrated the post-infection treatment efficacy of BXM against HPAIV infection in ducks, as it markedly decreased mortality with a significant reduction in virus shedding. Due to their slower progression of clinical symptoms, the duck model is more suitable than the chicken model for evaluating the efficacy of post-infection administration of BXM.

An improvement in the survival rate against HPAIV challenge among ducks administered at 0.5 mg/kg of BXM was confirmed. Moreover, all the birds survived at the dose of 2.5 mg/kg, confirming a dose-dependent effect on the clinical onset in delayed administration. On the other hand, one bird in the 12.5 mg/kg dose group developed leg paralysis. Previous studies indicated that viruses were detected in duck brains at 2–3 days post-HPAIV infection and cause neurological signs (Kishida et al., 2005; Spackman et al., 2023), implying that viruses might have invaded the duck’s nervous system before starting the treatment in this study. Neural tissue damage is known to be irreversible (Hosseini et al., 2018), thus delayed treatment reduces the probability of complete clinical recovery. Therefore, it is necessary to administer BXM soon after confirming virus infection in captive birds in situations of high risk of HPAIV infection.

The titers of virus recovered from oral swabs of ducks administered at 0.5 mg/kg significantly decreased from 2 dpid (*p*=0.012). Virus titers in oral swabs were below the detection limit from 4 dpid. A significant reduction in virus titer in oral swab was observed in the 12.5 and 2.5 mg/kg groups (*p*=0.001, *p*<0.001) from 1 dpid, and the titer was below the detection limit from 3 dpid. These findings suggest that administration at doses of 2.5 mg/kg or higher rapidly suppresses virus shedding. In addition, oral swabs showed a higher virus titer than cloacal swabs, indicating that oral swabs may be more appropriate for evaluating the inhibitory effect of BXM on virus shedding.

In the 12.5 mg/kg group, virus shedding was suppressed at 1 dpid, whereas virus shedding was recovered at 2 dpid, suggesting the potential involvement of drug-resistant viruses. Previous studies also demonstrated that drug-resistant viruses were promptly selected in the case of delayed administration (Kiso et al., 2025). The BXM-resistant variant of H5 HPAIV harboring the PA/I38T substitution, the most frequently emerging mutation under BXM treatment, exhibited an up to 48-fold increase in the 90% effective concentration (EC_90_) compared with the susceptible variant (Taniguchi et al., 2024), corresponding to the BXA concentration of 25.8 ng/mL. The concentration at 2 hours post-fifth administration transiently exceeded this value, suggesting that BXM could exert antiviral activity against resistant viruses when administered at more than 2.5 mg/kg. In this study, administration of BXM at a sufficient dose for five consecutive days led to a marked reduction in viral shedding in infected ducks. These results highlight the importance of predefining both the dose and duration of administration in the target avian species and adhering strictly to the established treatment protocol. Such optimization is important for considering and managing the potential risk of selecting resistant variants during the treatment of rare birds infected with HPAIV.

In this study, viral genome detection was performed by qRT-PCR in parallel with the monitoring of infectious virus shedding. The persistence of viral genome detection, even after infectious virus shedding disappeared, is consistent with previous reports of long-term persistence of non-infectious viral genome shedding in ducks (Wibawa et al., 2014). Therefore, while qRT-PCR is effective for detecting early infection, it may not be appropriate for determining the therapeutic efficacy of BXM.

Pharmacokinetic analysis confirmed that BXA was induced in plasma after oral administration of BXM, and its concentration increased in a dose-dependent manner. However, a decrease in the plasma concentration of BXA was confirmed after five days of BXM administration, probably due to the induction of metabolic enzymes that accelerate drug clearance. Consequently, this reduction in drug concentration does not support continuous administration of BXM as a prophylactic measure during the HPAIV endemic season. A previous study suggested C_24hr_, the plasma concentration of BXA at 24 hours post-administration, as a surrogate marker for the antiviral effect of the BXM using a mouse model of infection with the A/WSN/33 (H1N1) strain (Ando et al., 2020). However, this parameter was derived from experiments using a mouse-adapted neurotropic strain and may not be directly applicable to HPAIV infection in the avian model, which typically results in systemic infection. In the mouse model, C_24hr_ correlated strongly with the reduction of virus excretion rather than AUC_0–24hr_ and C_max_, indicating that maintaining effective concentrations was important. However, this study found that AUC_0–24hr_ and C_max_ showed a higher correlation with the suppression of virus shedding than C_24hr_, indicating that higher exposure to BXA may correlate with anti-HPAIV efficacy in birds. Although the results were obtained from a limited number of samples, these parameters may be more appropriate indicators for predicting treatment efficacy of BXM against HPAIV infection in birds.

## 5. Conclusion

This study highlights the potential of BXM as a post-infection treatment against HPAIV infection in birds, identifying a 48-hour therapeutic time window and an effective dose of 2.5 mg/kg for reducing mortality and virus shedding. The efficacy of post-infection administration could be associated with the pharmacokinetics of BXA in plasma, particularly AUC_0–24hr_ and C_max_, which can serve as effective surrogate markers. Comparing the pharmacokinetics of BXA in ducks with those in rare birds may facilitate establishment of effective BXM treatment protocols against HPAIV for each avian species. Although BXM administration to ducks infected with HPAIV may have resulted in the emergence of resistant viruses, five consecutive days of BXM administration suppressed virus shedding. Collectively, these findings provide fundamental data to inform rational administration design of BXM for the treatment of HPAIV infection in rare bird species, ultimately contributing to conservation efforts.

## CRediT authorship contribution statement

**Yo Shimazu:** Conceptualization, Methodology, Formal analysis, Investigation, Data Curation, Writing – Original Draft, Visualization, Funding acquisition**. Norikazu Isoda:** Conceptualization, Methodology, Validation, Investigation, Writing – Review & Editing. **Takahiro Hiono:** Conceptualization, Methodology, Validation, Investigation, Writing – Review & Editing. **Hew Yik Lim:** Investigation**. Daiki Kobayashi:** Investigation**. Misato Shibazaki:** Investigation. **Nguyen Bao Linh:** Investigation. **Tatsuru Morita:** Investigation. **Ryo Daniel Obara:** Conceptualization, Methodology, Writing – Review & Editing. **Mariko Miki:** Conceptualization, Methodology, Writing – Review & Editing. **Hiromi Osaki:** Conceptualization, Methodology, Validation, Writing – Review & Editing. **Takao Shishido:** Conceptualization, Methodology, Writing – Review & Editing. **Yoshinori Ikenaka:** Methodology, Validation, Formal analysis, Resources, Data Curation, Writing – Review & Editing, Supervision. **Yoshihiro Sakoda:** Conceptualization, Methodology, Validation, Resources, Writing – Review & Editing, Supervision, Project administration, Funding acquisition.

## Acknowledgments

We would like to express our sincere gratitude to Dr. Keita Matsuno (International Collaboration Unit, International Institute for Zoonosis Control, Hokkaido University, Japan) for his valuable advice on the pharmacokinetic/pharmacodynamic analysis. We also thank Ms. Mayumi Endo (Laboratory of Microbiology, Hokkaido University, Japan), Ms. Mai Tamba (One Health Research Center, Hokkaido University, Japan) and Dr. Yared Beyene (Laboratory of Toxicology, Hokkaido University, Japan) for their technical support.

## Data availability statements

All data are available within the manuscript.

---

Figure S1. (A) Survival rate and (B) mean clinical score of the ducks infected with Cr/Hok/22, Bs/Aki/16/, and Ws/Hok/11.

Table S1. Clinical symptoms and HI titers in serum of ducks infected with Cr/Hok/22 followed by the administration of BXM at 12.5, 2.5, 0.5, or 0 mg/kg.

Table S2. Virus titer (upper) and Cp value (lower) of oral and cloacal swabs from the ducks infected with Cr/Hok/22 followed by the administration of BXM at 12.5, 2.5, 0.5, or 0 mg/kg.

Table S3. Plasma concentration of BXA after the first and fifth administration of BXM at 12.5 and 2.5 mg/kg.

## References

Alexander, D.J., Brown, I.H., 2009. History of highly pathogenic avian influenza. Rev. Sci. Tech. l’OIE 28, 19–38. 10.20506/rst.28.1.1856

Ando, Y., Noshi, T., Sato, K., Ishibashi, T., Yoshida, Y., Hasegawa, T., Onishi, M., Kitano, M., Oka, R., Kawai, M., Yoshida, R., Sato, A., Shishido, T., Naito, A., 2020. Pharmacokinetic and pharmacodynamic analysis of baloxavir marboxil, a novel cap-dependent endonuclease inhibitor, in a murine model of influenza virus infection. J. Antimicrob. Chemother. 76, 189–198. 10.1093/jac/dkaa393

Banyard, A.C., Bennison, A., Byrne, A.M.P., Reid, S.M., Lynton-Jenkins, J.G., Mollett, B., Silva, D.D., Peers-Dent, J., Finlayson, K., Hall, R., Blockley, F., Blyth, M., Falchieri, M., Fowler, Z., Fitzcharles, E.M., Brown, I.H., James, J., 2024. Detection and spread of high pathogenicity avian influenza virus H5N1 in the Antarctic Region. Nat. Commun. 15, 7433. 10.1038/s41467-024-51490-8

Bertelsen, M.F., Klausen, J., Holm, E., Grøndahl, C., Jørgensen, P.H., 2007. Serological response to vaccination against avian influenza in zoo-birds using an inactivated H5N9 vaccine. Vaccine 25, 4345–4349. 10.1016/j.vaccine.2007.03.043

Bordes, L., Germeraad, E.A., Roose, M., Eijk, N.M.H.A. van, Engelsma, M., Poel, W.H.M. van der, Vreman, S., Beerens, N., 2024. Experimental infection of chickens, Pekin ducks, Eurasian wigeons and Barnacle geese with two recent highly pathogenic avian influenza H5N1 clade 2.3.4.4b viruses. Emerg. Microbes Infect. 13, 2399970. 10.1080/22221751.2024.2399970

Checkmahomed, L., Padey, B., Pizzorno, A., Terrier, O., Rosa-Calatrava, M., Abed, Y., Baz, M., Boivin, G., 2020. In vitro combinations of baloxavir acid and other inhibitors against seasonal influenza A viruses. Viruses 12, 1139. 10.3390/v12101139

Dias, A., Bouvier, D., Crépin, T., McCarthy, A.A., Hart, D.J., Baudin, F., Cusack, S., Ruigrok, R.W.H., 2009. The cap-snatching endonuclease of influenza virus polymerase resides in the PA subunit. Nature 458, 914–918. 10.1038/nature07745

Fujimoto, Y., Ogasawara, K., Isoda, N., Hatai, H., Okuya, K., Watanabe, Y., Takada, A., Sakoda, Y., Saito, K., Ozawa, M., 2022. Experimental and natural infections of white-tailed sea eagles (Haliaeetus albicilla) with high pathogenicity avian influenza virus of H5 subtype. Front. Microbiol. 13, 1007350. 10.3389/fmicb.2022.1007350

Heine, H.G., Foord, A.J., Wang, J., Valdeter, S., Walker, S., Morrissy, C., Wong, F.Y., Meehan, B., 2015. Detection of highly pathogenic zoonotic influenza virus H5N6 by reverse-transcriptase quantitative polymerase chain reaction. Virol. J. 12, 18. 10.1186/s12985-015-0250-3

Hiono, T., Okamatsu, M., Yamamoto, N., Ogasawara, K., Endo, M., Kuribayashi, S., Shichinohe, S., Motohashi, Y., Chu, D.-H., Suzuki, M., Ichikawa, T., Nishi, T., Abe, Y., Matsuno, K., Tanaka, K., Tanigawa, T., Kida, H., Sakoda, Y., 2016. Experimental infection of highly and low pathogenic avian influenza viruses to chickens, ducks, tree sparrows, jungle crows, and black rats for the evaluation of their roles in virus transmission. Vet. Microbiol. 182, 108–115. 10.1016/j.vetmic.2015.11.009

Hosseini, S., Wilk, E., Michaelsen-Preusse, K., Gerhauser, I., Baumgärtner, W., Geffers, R., Schughart, K., Korte, M., 2018. Long-term neuroinflammation induced by influenza A virus infection and the impact on hippocampal neuron morphology and function. J. Neurosci. 38, 3060–3080. 10.1523/jneurosci.1740-17.2018

Isoda, N., Onuma, M., Hiono, T., Sobolev, I., Lim, H.Y., Nabeshima, K., Honjyo, H., Yokoyama, M., Shestopalov, A., Sakoda, Y., 2022. Detection of new H5N1 high pathogenicity avian influenza viruses in winter 2021–2022 in the far east, which are genetically close to those in Europe. Viruses 14, 2168. 10.3390/v14102168

Itzstein, M. von, Wu, W.-Y., Kok, G.B., Pegg, M.S., Dyason, J.C., Jin, B., Phan, T.V., Smythe, M.L., White, H.F., Oliver, S.W., Colman, P.M., Varghese, J.N., Ryan, D.M., Woods, J.M., Bethell, R.C., Hotham, V.J., Cameron, J.M., Penn, C.R., 1993. Rational design of potent sialidase-based inhibitors of influenza virus replication. Nature 363, 418–423. 10.1038/363418a0

Katzner, T.E., Blackford, A.V., Donahue, M., Gibbs, S.E.J., Lenoch, J., Martin, M., Rocke, T.E., Root, J.J., Styles, D., Cooper, S., Dean, K., Dvornicky-Raymond, Z., Keller, D., Sanchez, C., Dunlap, B., Grier, T., Jones, M.P., Nitzel, G., Patrick, E., Purcell, M., Specht, A.J., Suarez, D.L., 2025. Safety and Immunogenicity of Poultry Vaccine for Protecting Critically Endangered Avian Species against Highly Pathogenic Avian Influenza Virus, United States. Emerg. Infect. Dis. 31, 1131–1139. 10.3201/eid3106.241558

Kishida, N., Sakoda, Y., Isoda, N., Matsuda, K., Eto, M., Sunaga, Y., Umemura, T., Kida, H., 2005. Pathogenicity of H5 influenza viruses for ducks. Arch. Virol. 150, 1383–1392. 10.1007/s00705-004-0473-x

Kiso, M., Uraki, R., Yamayoshi, S., Kawaoka, Y., 2025. Efficacy of baloxavir marboxil against bovine H5N1 virus in mice. Nat. Commun. 16, 5356. 10.1038/s41467-025-60791-5

Liu, S., Zhuang, Q., Wang, S., Jiang, W., Jin, J., Peng, C., Hou, G., Li, J., Yu, J., Yu, X., Liu, H., Sun, S., Yuan, L., Chen, J., 2020. Control of avian influenza in China: Strategies and lessons. Transbound. Emerg. Dis. 67, 1463– 1471. 10.1111/tbed.13515

Meijer, A., Goot, J.A. van der, Koch, G., Boven, M. van, Kimman, T.G., 2004. Oseltamivir reduces transmission, morbidity, and mortality of highly pathogenic avian influenza in chickens. Int. Congr. Ser. 1263, 495–498. 10.1016/j.ics.2004.01.020

Oh, S., Martelli, P., Hock, O.S., Luz, S., Furley, C., Chiek, E.J., Wee, L.C., Keun, N.M., 2005. Field study on the use of inactivated H5N2 vaccine in avian species. Vet. Rec. 157, 299–300. 10.1136/vr.157.10.299

Okamatsu, M., Ozawa, M., Soda, K., Takakuwa, H., Haga, A., Hiono, T., Matsuu, A., Uchida, Y., Iwata, R., Matsuno, K., Kuwahara, M., Yabuta, T., Usui, T., Ito, H., Onuma, M., Sakoda, Y., Saito, T., Otsuki, K., Ito, T., Kida, H., 2017. Characterization of highly pathogenic avian influenza virus A(H5N6), Japan, November 2016. Emerg. Infect. Dis. 23, 691–695. 10.3201/eid2304.161957

Philippa, J.D.W., Munster, V.J., Bolhuis, H. van, Bestebroer, T.M., Schaftenaar, W., Beyer, W.E.P., Fouchier, R.A.M., Kuiken, T., Osterhaus, A.D.M.E., 2005. Highly pathogenic avian influenza (H7N7): Vaccination of zoo birds and transmission to non-poultry species. Vaccine 23, 5743–5750. 10.1016/j.vaccine.2005.09.013

Sacristán, C., Ewbank, A.C., Porras, P.I., Pérez-Ramírez, E., Torre, A. de la, Briones, V., Iglesias, I., 2024. Novel epidemiologic features of high pathogenicity avian influenza virus A H5N1 2.3.3.4b panzootic: a review. Transbound. Emerg. Dis. 2024, 5322378. 10.1155/2024/5322378

Sakoda, Y., Ito, H., Uchida, Y., Okamatsu, M., Yamamoto, N., Soda, K., Nomura, N., Kuribayashi, S., Shichinohe, S., Sunden, Y., Umemura, T., Usui, T., Ozaki, H., Yamaguchi, T., Murase, T., Ito, T., Saito, T., Takada, A., Kida, H., 2012. Reintroduction of H5N1 highly pathogenic avian influenza virus by migratory water birds, causing poultry outbreaks in the 2010–2011 winter season in Japan. J. Gen. Virol. 93, 541–550. 10.1099/vir.0.037572-0

Schlachter, A.-L.D., Furman, N., Byrne, A.M.P., Reid, S.M., Smith, S.J., Maskell, D., Mollett, B.C., Peers-Dent, J., Falchieri, M., Schock, A., Banyard, A.C., Brown, I.H., Núñez, A., Manuscript, A.-L.D.S.A.N.F.H.E.C.T.T.S.W.A.T.P.O.T., 2025. High pathogenicity avian influenza H5N1 clade 2.3.4.4b natural infection in captive Humboldt penguins (Spheniscus humboldti). Avian Pathol. : J. WVPA 1–21. 10.1080/03079457.2025.2513338

Soda, K., Tomioka, Y., Usui, T., Uno, Y., Nagai, Y., Ito, H., Hiono, T., Tamura, T., Okamatsu, M., Kajihara, M., Nao, N., Sakoda, Y., Takada, A., Ito, T., 2022. Susceptibility of herons (family: Ardeidae) to clade 2.3.2.1 H5N1 subtype high pathogenicity avian influenza virus. Avian Pathol. 51, 146–153. 10.1080/03079457.2021.2022599

Spackman, E., Pantin-Jackwood, M.J., Lee, S.A., Prosser, D., 2023. The pathogenesis of a 2022 North American highly pathogenic clade 2.3.4.4b H5N1 avian influenza virus in mallards (Anas platyrhynchos). Avian Pathol. 52, 219–228. 10.1080/03079457.2023.2196258

Spickler, A.R., Trampel, D.W., Roth, J.A., 2008. The onset of virus shedding and clinical signs in chickens infected with high-pathogenicity and low-pathogenicity avian influenza viruses. Avian Pathol. 37, 555–577. 10.1080/03079450802499118

Taniguchi, K., Noshi, T., Omoto, S., Sato, A., Shishido, T., Matsuno, K., Okamatsu, M., Krauss, S., Webby, R.J., Sakoda, Y., Kida, H., 2024. The impact of PA/I38 substitutions and PA polymorphisms on the susceptibility of zoonotic influenza A viruses to baloxavir. Arch. Virol. 169, 29. 10.1007/s00705-023-05958-5

Twabela, A., Okamatsu, M., Matsuno, K., Isoda, N., Sakoda, Y., 2020. Evaluation of baloxavir marboxil and peramivir for the treatment of high pathogenicity avian influenza in chickens. Viruses 12, 1407. 10.3390/v12121407

Usui, T., Soda, K., Sumi, K., Ozaki, H., Tomioka, Y., Ito, H., Murase, T., Kawamoto, T., Miura, M., Komatsu, M., Imanishi, T., Kurobe, M., Ito, T., Yamaguchi, T., 2020. Outbreaks of highly pathogenic avian influenza in zoo birds caused by HA clade 2.3.4.4 H5N6 subtype viruses in Japan in winter 2016. Transbound. Emerg. Dis. 67, 686–697. 10.1111/tbed.13386

Vergara-Alert, J., Fernández-Bellon, H., Busquets, N., Alcántara, G., Delclaux, M., Pizarro, B., Sánchez, C., Sánchez, A., Majó, N., Darji, A., 2011. Comprehensive Serological Analysis of Two Successive Heterologous Vaccines against H5N1 Avian Influenza Virus in Exotic Birds in Zoos. Clin. Vaccine Immunol. 18, 697–706. 10.1128/cvi.00013-11

Wibawa, H., Bingham, J., Nuradji, H., Lowther, S., Payne, J., Harper, J., Junaidi, A., Middleton, D., Meers, J., 2014. Experimentally infected domestic ducks show efficient transmission of indonesian H5N1 highly pathogenic avian influenza Vvirus, but lack persistent viral shedding. PLoS ONE 9, e83417. 10.1371/journal.pone.0083417

WOAH, 2024. Avian influenza (Including infection with high pathogenicity avian influenza virus), in: WOAH Terrestrial Manual 2025.

Xie, Z., Yang, J., Jiao, W., Li, X., Iqbal, M., Liao, M., Dai, M., 2025. Clade 2.3.4.4b highly pathogenic avian influenza H5N1 viruses: knowns, unknowns, and challenges. J. Virol. e0042425. 10.1128/jvi.00424-25

