## Supplementary material for "Evaluation of therapeutic effect of baloxavir marboxil against high pathogenicity avian influenza virus infection in duck model": bioRxiv_Duck_SupFig.pdf

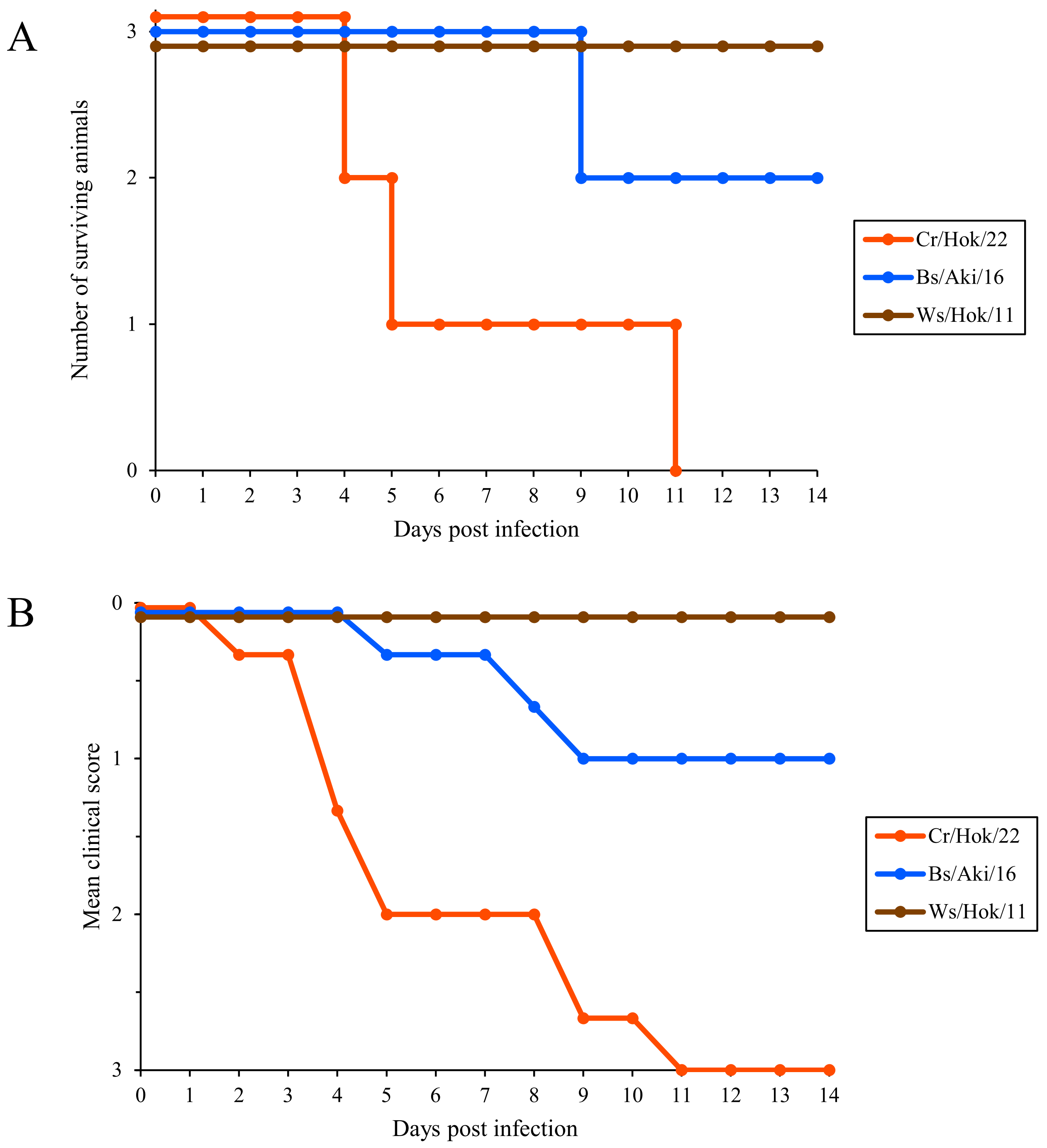


Figure S1. (A) Survival rate and (B) mean clinical score of the ducks infected with Cr/Hok/22, Bs/Aki/16/, and Ws/Hok/11.
